# Conditional-longitudinal brain growth charts detect MRI changes with birth weight and psychopathology

**DOI:** 10.64898/2025.12.24.696365

**Authors:** Eren Kafadar, Margaret Gardner, Lena Dorfschmidt, Jonathan A. Berken, Audrey C. Luo, Kevin Y. Sun, Richard A.I. Bethlehem, Sara B. DeMauro, Ran Barzilay, Varun Warrier, Tyler M. Moore, Jakob Seidlitz, Heather H. Burris, Theodore D. Satterthwaite, Russell T. Shinohara, Aaron F. Alexander-Bloch

**Affiliations:** Department of Psychiatry, Perelman School of Medicine, University of Pennsylvania, Philadelphia, PA; Department of Child and Adolescent Psychiatry and Behavioral Science, The Children’s Hospital of Philadelphia, Philadelphia, PA; Lifespan Brain Institute of CHOP and Penn, Philadelphia, PA; Department of Pediatrics, Perelman School of Medicine at the University of Pennsylvania, Philadelphia, PA, USA; Division of Neonatology, Department of Pediatrics, The Children’s Hospital of Philadelphia, Philadelphia, PA, USA; Division of Gastroenterology, Hepatology, and Nutrition, Department of Pediatrics, The Children’s Hospital of Philadelphia, Philadelphia, PA, USA; Penn Lifespan Informatics and Neuroimaging Center (PennLINC), Department of Psychiatry, Perelman School of Medicine, University of Pennsylvania, Philadelphia, PA, USA; Department of Psychology, University of Cambridge, Cambridge, UK; Department of Psychiatry, University of Cambridge, Cambridge, UK; Leonard Davis Institute of Health Economics, University of Pennsylvania, Philadelphia, Pennsylvania; Department of Biostatistics, Epidemiology and Informatics, University of Pennsylvania, Philadelphia, PA; Department of Obstetrics and Gynecology, Perelman School of Medicine at the University of Pennsylvania, Philadelphia, PA; Penn Statistics in Imaging and Visualization Center, Department of Biostatistics, Epidemiology, and Informatics, University of Pennsylvania, Philadelphia, PA 19104; Center for AI and Data Science for Integrated Diagnostics, University of Pennsylvania, Philadelphia, PA 19104

## Abstract

**Importance:** Brain maturation varies between individuals, particularly during dynamic developmental periods like adolescence. Directly assessing differences in longitudinal trajectories can reveal deviations from normative patterns.

**Objective:** We present novel conditional-longitudinal normative models that characterize variability in brain maturation. We utilize these models to examine whether differences in longitudinal trajectories are associated with birth weight (BW), gestational age (GA), and longitudinal psychopathology derived from behavioral assessments.

**Design:** Cross-sectional and conditional-longitudinal normative models were developed for brain volumes derived from the first two neuroimaging timepoints from the Adolescent Brain Cognitive Development (ABCD) Study. Conditional-longitudinal models index an individual’s expected brain volume at follow-up conditioned on their baseline measurement. Models were fit with split-half cross-validation on demographically matched samples.

**Setting:** The ABCD Study is a multi-site, population-based study

**Participants:** Participants were excluded based on imaging quality flags and missing data, leaving 10,830 at baseline and 7,262 at follow-up.

**Exposures:** BW and GA were derived from parent-report questionnaires. General psychopathology scores were calculated using a bifactor model.

**Main Outcomes and Measures:** We calculated cross-sectional and conditional-longitudinal centiles, respectively quantifying individual deviations in size and change between timepoints. Sensitivity analyses included covariates for parental income and education as well as current weight and height.

**Results:** The sample was 10,830 at baseline (48.2% F,age 9-10y) and 7,262 at follow-up (46.6% F,age 11-13y). Conditional-longitudinal centiles were sensitive to individual differences in brain change between timepoints. Lower BW was associated with lower conditional-longitudinal centiles, suggesting larger decreases in brain volumes over time (27 regions p_fdr_<0.05, β_max_=0.08). Lower conditional-longitudinal centiles were associated with greater increases in psychopathology scores, suggesting with increased psychopathology brain volumes show greater decrease (37 regions p_fdr_<0.05, β_max_=0.06). Notably, changes in psychopathology were not related to brain size at either timepoint, indexed by cross-sectional centiles.

**Conclusions and Relevance:** Models that capture individual-level deviations from expected growth trajectories, rather than static positions on a growth curve, are particularly informative for assessing developmental change. Novel conditional-longitudinal models address this gap in lifespan brain imaging. Using this framework, we demonstrate robust associations between individual trajectory deviations, perinatal adversity, and longitudinally assessed mental health symptoms. Condition-longitudinal models hold promise for applications across psychiatric neuroscience, from development to aging.

**Key Points:** 

**Question:** How do differences in brain maturation trajectories, quantified by novel conditional-longitudinal models, relate to perinatal factors and mental health in adolescence?

**Findings:** In this longitudinal analysis of neuroimaging data from the Adolescent Brain Cognitive Development (ABCD) Study, conditional-longitudinal normative models revealed that trajectories of brain maturation in adolescence are associated with birth weight, and with longitudinal changes in mental health.

**Meaning:** Conditional longitudinal models detect inter-individual variability in brain maturation, which is related to both perinatal factors and concurrent changes in psychopathology.

## Introduction

Normative models of neuroimaging data are increasingly deployed to benchmark inter-individual differences in quantitative features against a population reference model. Similar to pediatric growth charts of height or weight, such “brain charts,” can boost interpretability and signal-to-noise ratio to detect clinical differences.^1–3^ Previous normative modeling studies largely model longitudinal data cross-sectionally, which does not directly address variability in longitudinal change.^4–6^ Over time, individuals with extreme centiles tend to move closer to the mean, suggesting different expected growth velocities based on initial position on the growth chart.^7,8^ Longitudinal normative models can account for this regression-to-the-mean by benchmarking change between timepoints, which may be particularly useful in highly dynamic periods of brain development such as adolescence. Here, we present a conditional-longitudinal modeling framework in which within-individual change is quantified by conditioning follow-up brain volume measurements on baseline values. Harnessing longitudinal anatomical brain MRI data from the Adolescent Brain Cognitive Development (ABCD) Study,^9^ we use this approach to test for brain trajectory differences associated with perinatal developmental factors (e.g., birth weight) and with contemporaneous measures of psychopathology.

Extensive research links perinatal factors including birth weight (BW) and gestational age (GA) to brain MRI differences that persist into adolescence and adulthood. In general, prematurity and low birth weight have been associated with smaller regional brain volumes.^10–14^ However, few studies provide a longitudinal perspective to assess whether effects of perinatal adversity are stable throughout the lifespan, or whether they continue to exert influence on the dynamics of brain maturation. Studies have suggested that prematurity may result in slower brain growth in childhood,^15,16^ and there may be age-varying relationships between birth weight and brain volume in adolescence.^17,18^ However, prior work has relied on modeling approaches that do not fully disentangle static differences from differences in how the brain changes. For instance, if birth weight is correlated with brain volume at baseline, and individuals with larger brains tend to have larger absolute change between timepoints, then associations with “change” might simply reflect a propagation of baseline differences rather than meaningful differences in longitudinal trajectories. Conditional-longitudinal models resolve this uncertainty by quantifying the degree of expected change for each individual based on their baseline brain volumes. This approach uncovers whether smaller brains associated with perinatal adversity continue to mature similarly to smaller brains without perinatal adversity, or whether the effects of perinatal adversity may be amplified as the brain continues to mature.

In addition to an association with perinatal factors, individual deviations from normative growth patterns may be related to changes in mental health.^19^ Adolescence is a critical period for both brain development and sensitivity to mental illness.^20,21^ As such, understanding the relationship between brain maturation and mental health trajectories is a critical goal. Baseline neuroimaging features in ABCD have previously been linked to overall psychopathology, indexed by the “P-factor”. ^22,23^ Smaller studies have also suggested that structural brain changes can track with severity of psychiatric symptoms over time.^24,25^ However, the potential utility of longitudinal normative models remains untested. Here, we introduce conditional-longitudinal models to address this gap, testing whether within-person changes in P-factor are associated with deviations from expected longitudinal brain development benchmarked by our models.

## Methods

To ensure reproducibility and maximize useability of conditional-longitudinal normative models for other researchers, all analyses were run on publicly available tabular ABCD Data Release 5.1(https://www.nbdc-datahub.org/).^26^All analyses were run in R v4.2.3, and code is available at https://github.com/BGDlab/longitudinal-models-abcd

### Participants

Questionnaire data and brain volumes derived from anatomical MRI scans at ABCD t_1_ (baseline) and t_2_ (approximately 2 years a later) spanned ages 9-13 years.^9^ Caregivers of participants provided informed consent and study sites received IRB approval. Exclusions were based on data availability and image quality flags (**Supplemental Materials Section 1 (S1)**).

### Variables of Interest

#### Anatomical MRI

Brain regions investigated included cortical volumes (Desikan-Killiany parcellation)^27^ and subcortical volumes (‘Aseg’ parcellation).^28^ Global tissue volumes included total cortical gray matter, total cerebral white matter, total ventricles, total subcortical gray matter. Intracranial volume was utilized in sensitivity analyses. Results reported in the main text pertain to the regional volumes; findings for global tissue volumes are found in **Supplemental Tables 6-8**. Details on MRI image acquisition and processing are described in REF^9^ and **Supplement S2**.^9^

#### Birth-Related Factors

Parent-report questionnaires provided gestational age (weeks), birth weight (ounces), and neonatal complications (Supplementary Table 1). Given their high co-linearity, birth weight was corrected for gestational age using the *growthstandards* package v0.1.6.^29^ This package uses Intergrowth standards growthcharts,^30^ or birth weight percentiles.

#### Birth weight PGS

Birth weight polygenic scores (PGS-BW) were calculated using PRS-CS^31^ trained on GWAS^32^ from the Early Growth Genetics^33^ consortium, limited to the European-ancestry subsample (**Supplement S3**).

#### Psychopathology

P-factor measured general psychopathology derived from a series of mental health questionnaires completed at t_1_ and t_2_. A longitudinal bifactor model was used where each mental health interview item loads onto the general factor and one of multiple orthogonal sub-factors. Analyses were based on the split-sample exploratory-to-confirmatory procedure described in Moore et al 2020 (**Supplement S4**).^22^

### Normative Models

Using the *gamlss* v5.4.22 package,^34^ we fit cross-sectional normative models to t_1_ or t_2_, respectively and conditional-longitudinal models to t_1_ and t_2_ together. The model selection space included age and sex as covariates. A random effect for site was used in all models to harmonize differences in location and scale across sites^1^. Conditional-longitudinal models of t_2_ brain volumes included t_1_ volumes for conditioning on baseline measurements and the interscan interval as an optional covariate in the model selection space, yielding participant-specific reference distributions for t_2_ brain volumes given t_1_ brain volumes (**Supplement S5**).

Normative models were fit in two matched split-halves based on the ABCD Reproducible Matched Samples.^35,36^ To prevent data leakage, centiles (i.e., participant-specific deviation from normative models) for split-half A were calculated using model estimates from split-half B, and vice versa. Optional covariates included in the models were comparable between split-halves (**Table 1**; **Supplementary Table 2**), and final model parameters can be found in **Supplementary Tables 3-5**.

**Table 1.**
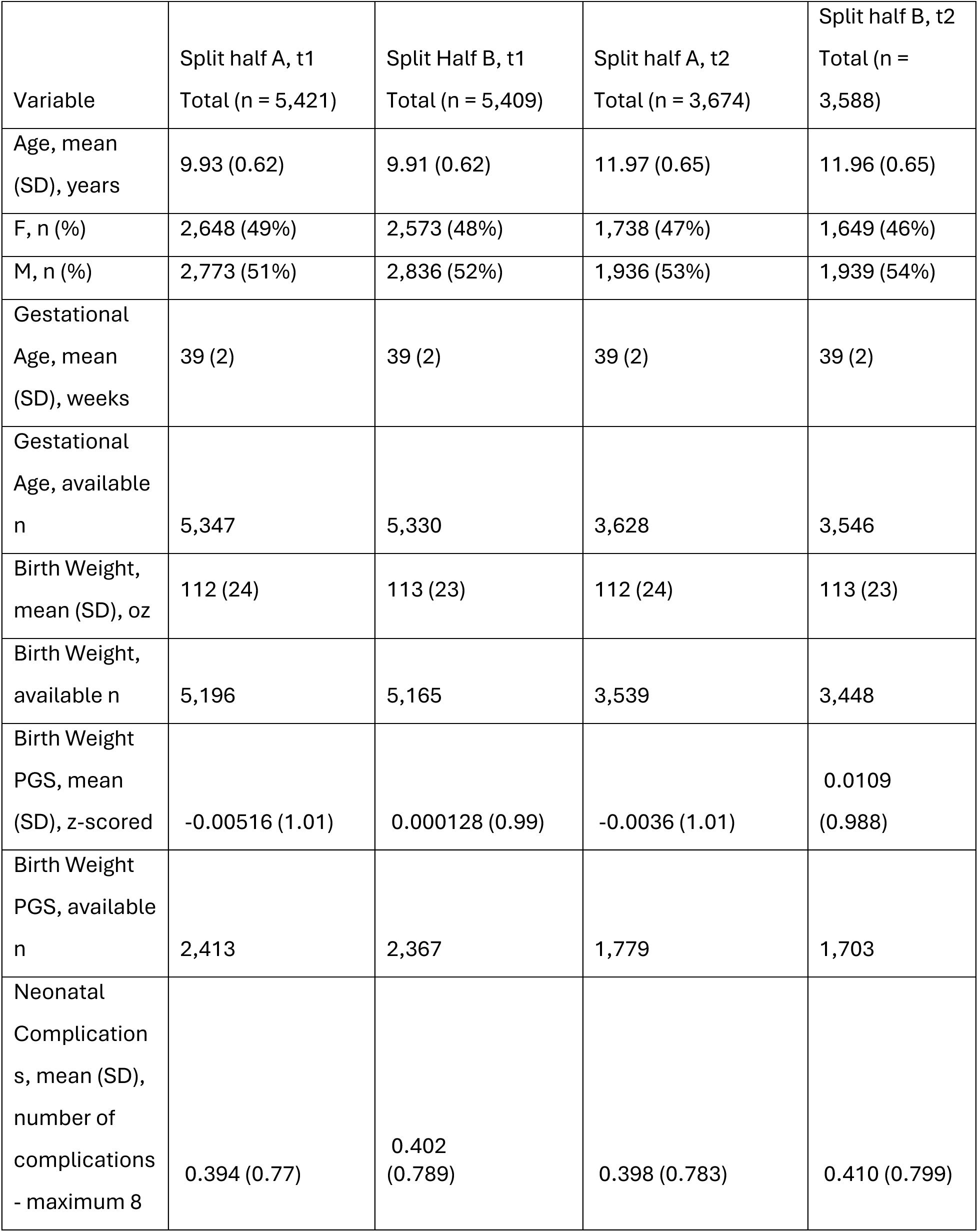
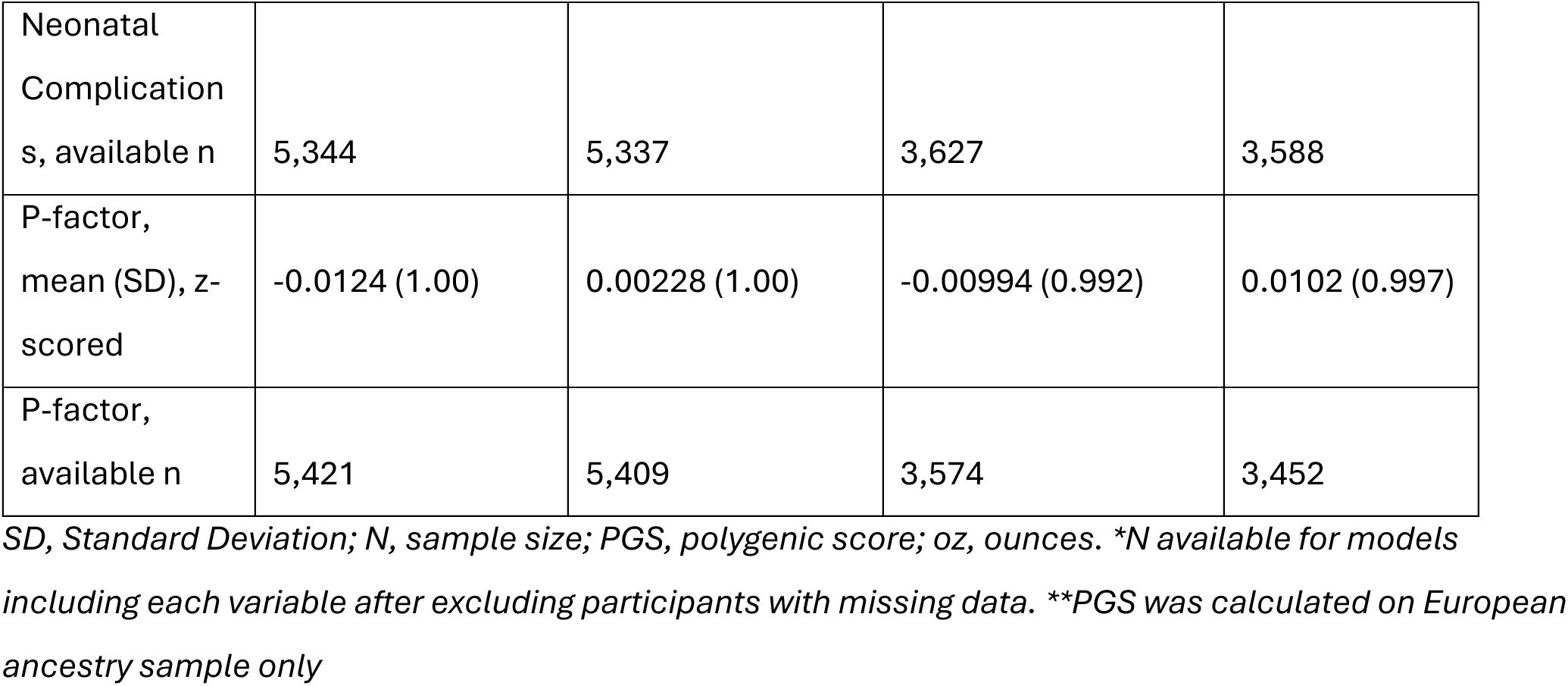
Distribution of modeling variables and variables of interest for each split sample and timepoint.

### Statistical Analyses

Linear regression tested the associations between centiles and other variables of interest. To normalize variables and enable comparison of effects, all variables of interest were z-scored, and centiles were converted into “standardized deviation scores” using the *qnorm* function in R (equivalent to z-scores for normally distributed data). To generate a summary value across regional centiles, a cumulative deviation score, the centile Mahalanobis distance, was calculated (CMD^1^;**Supplement S5**). To account for differences in baseline psychopathology scores, baseline P-factor scores was included as a covariate in regressions with change in P-factor scores. To account for genetic ancestry effects, the first ten ancestry principal components were included as covariates in regressions with PGS-BW. Multiple comparisons in P-values were corrected using the Benjamini-Hochberg procedure within four global tissue volumes, and within all regional volumes. Standardized βs with p_corrected_ < 0.05 are reported as significant. Sensitivity analyses included parental income and education, and current height and weight as covariates. To explore associations above and beyond brain size as a whole, regressions were run including an intracranial volume covariate (**Supplement S6**). All anatomical visualizations were rendered using *ggseg* v1.6.5.^37^

## Results

### Unique Information Provided by Conditional-longitudinal Normative Models

Conditional-longitudinal models computed at t_2_ were sensitive to individual brain change between timepoints, yielding distinct information from cross-sectional models computed at t_1_ or t_2_. As illustrated in **Figure 1**, because conditional-longitudinal models are conditioned on a given participant’s brain volumes measured at t_1,_ the resulting centile lines are highly participant-specific. Using CMD as a cumulative measure of deviation, there was a high degree of correlation between CMD_t1_ and CMD_t2_ cross-sectional centiles (CMD_t1-t2_,r = 0.81), while CMD_long_ centiles were correlated with cross-sectional centiles to a much lower degree (CMD_t1-long_,r= 0.19; CMD_t2-long_, r=0.20; **Supplementary Figure 1**).

**Figure 1.**
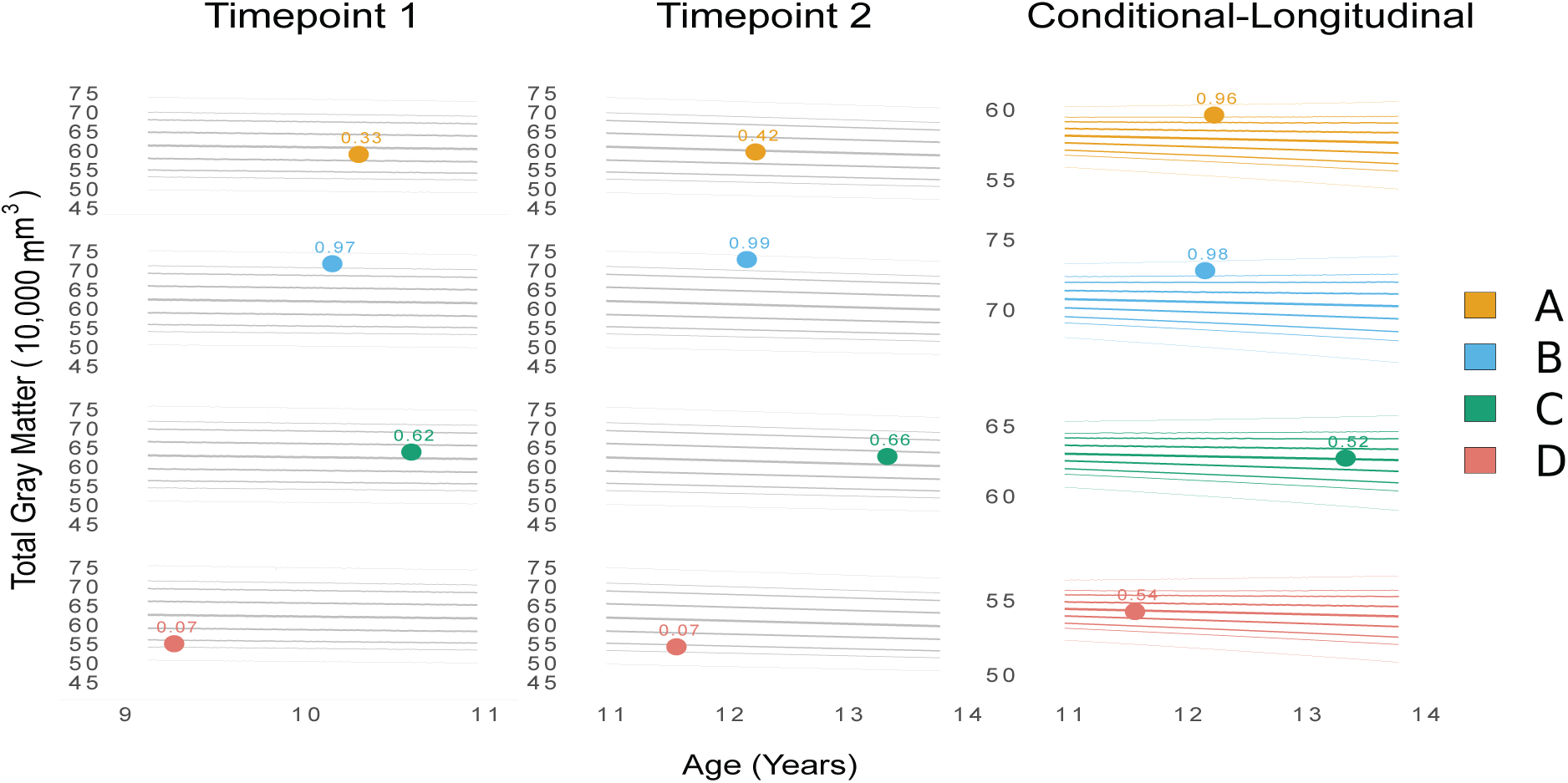
Conditional-Longitudinal Models Uniquely Characterize Differences in Trajectories. Centile lines from conditional-longitudinal normative models diverge across participants in contrast to centile lines from cross-sectional normative models, as shown for total gray matter volume of four illustrative ABCD participants. At baseline (t_1_), Participant A (yellow) and Participant C (green) were closer to the middle of the distribution modeled by the t_1_ cross-sectional normative model (centile_t1A_ = 0.33, centile_t1C_ = 0.62), while Participant B (blue) and Participant D (red) were closer to the tails of the distribution (centile_t1B_ = 0.97, centile_t1D_ = 0.07). At follow up (t_2_), when benchmarked against the t_2_ cross-sectional normative model, Participant A moved 9 centiles upwards while Participant B moved two centiles upward (centile_t2A_ = 0.42, centile_t2C_ = 0.99). However, when benchmarked against the conditional-longitudinal model, both Participants had relatively high centiles (centile_longA_ = 0.96, centile_longC_ = 0.98), reflecting relatively extreme changes in gray matter volume over time. On the other hand, when benchmarked against the cross-sectional models, Participant C moved four centiles upward from centile_t1C_ = 0.62 to centile_t2C_ = 0.66, but this change was close to the 50^th^ centile line estimated by the conditional-longitudinal model (centile_longC_ = 0.52). Participant D remained at the same relatively low centile when benchmarked against the cross-sectional models (centile_t1D_ = 0.07, centile_t2D_ = 0.07), and their t_2_ volume was also close to the 50^th^ centile line estimated by the conditional-longitudinal model (centile_longD_ = 0.54).

### Birth Weight Associated with Conditional-longitudinal Centiles

Next, we examined the associations between both cross-sectional and conditional-longitudinal centiles and perinatal factors. Smaller BW and younger GA were each associated with lower cross-sectional centiles at both timepoints, corresponding to smaller brain volumes (BW_t1_, N_regions_ (p_fdr_<0.05) = 96, β_range_= (0.023, 0.22); BW_t2_, N_regions_ (p_fdr_<0.05) = 94, β_range_= (−0.026, 0.22); GA t_1_, N_regions_ (p_fdr_<0.05) = 65, β_range_= (−0.045, 0.086); GA t_2_, N_regions_ (p_fdr_<0.05) = 37, β_range_= (−0.048, 0.075)). The strength of these associations was greater for BW and varied regionally (**Figure 2**). An exception was centiles for ventricles, which showed an inverse relationship such that ventricular volume was smaller given older GAs at birth.

**Figure 2.**
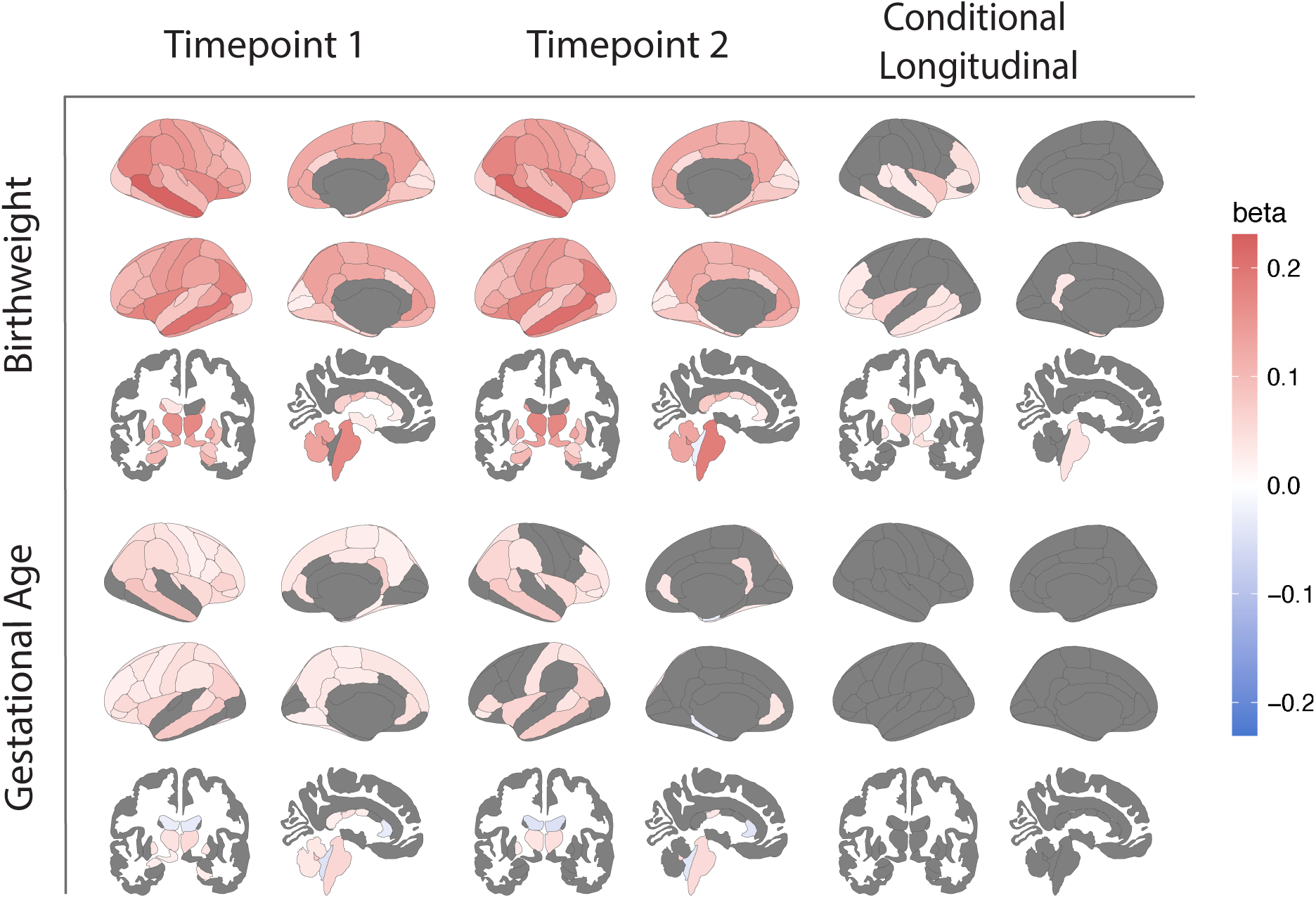
Lower Birth Weight is Related to Smaller Size and Smaller Longitudinal Centiles. Lower birth weight and smaller gestational age (GA) were associated with overall lower cross-sectional centiles (corresponding to smaller size), while only lower birth weight was associated with lower conditional-longitudinal centiles (corresponding to greater-than-expected decreases in volume). All results shown were statistically significant at FDR-corrected p_fdr_<0.05, regions that did not survive multiple comparisons are in gray. (Birth weight_t1_, N_regions_ (pfdr<0.05) = 96, standardized β_range_= (0.023, 0.22); birth weight_t2,_ N_regions_ (pfdr<0.05)= 94, *β*_range_= (−0.026, 0.22), birth weight_long_ N_regions_ (pfdr<0.05) = 27, β_range_= (0.030, 0.083); GA t_1,_ N_regions_ (pfdr<0.05) = 65, β_range_= (−0.045, 0.086); GA t_2_, N_regions_ (pfdr<0.05) = 37, β_range_= (−0.048, 0.075)).For detailed information on regression statistics across all regions and global tissue volumes see **Supplementary Tables 6-8**.

Testing the relationship between perinatal factors and brain change, we found that conditional-longitudinal centiles were positively associated with BW, and no significant associations were found with GA. Smaller BW was associated with smaller regional volumes at t_2_ after accounting for the size of regional volumes at t_1_, especially for frontal and temporal cortical volumes, bilateral thalamus, and brainstem volumes (N_regions_ (p_fdr_<0.05) = 27, β_range_= (0.030, 0.083)) (**Figure 2**).

Given the degree of covariance between BW and GA, we calculated a BW-percentile using published Intergrowth growth charts^29^. The BW-percentile provides a measure of BW after accounting for GA. The relationship between BW-percentile and brain volume centiles closely followed results from linear regressions with BW itself, showing robustness of BW – brain associations when the relationship between BW and GA is accounted for (BW-percentile _t1_, N_regions_ (p_fdr_<0.05) = 98, β_range_= (0.018, 0.16); BW-percentile _t2_, N_regions_ (p_fdr_<0.05) = 96, β_range_= (0.022, 0.17); BW-percentile _long_, N_regions_ (p_fdr_<0.05) = 31, β_range_= (0.023, 0.059), p_fdr_ <0.05) (**Supplementary Figure 2**). Effect sizes for BW-percentile were largely robust to including GA as a covariate (BW-percentile_t1_, N_regions_ (p_fdr_<0.05) = 98, β_range_= (0.018, 0.16); BW-percentile _t2_, N_regions_ (p_fdr_<0.05) = 96, β_range_= (0.022, 0.17); BW-percentile _long_, N_regions_ (p_fdr_<0.05) = 33, β_range_= (0.022, 0.061)) (**Supplementary Tables 6-8)**.

We performed additional analyses to understand underlying factors contributing to these observed relationships. First, we calculated a polygenic score for BW (PGS-BW) aggregating genetic factors associated with BW based on a recent genome-wide association study^31^ (**Supplement S3**). In support of the generalizability of PGS-BW to the European-ancestry subsample of ABCD (N_t1_ = 4780, N_t2_ = 3482), we confirmed a small but significant association between PGS-BW and the phenotype of BW in our sample (r = 0.14, p < 2.2×10^-16^). Higher PGS-BW was associated with higher brain centiles (PGS-BW_t1_, N_regions_ (p_fdr_<0.05) = 73, β_range_= (0.031, 0.082); PGS-BW_t2,_ N_regions_ (p_fdr_<0.05) = 71, β_range_= (0.035, 0.097, p_fdr_<0.05)). Moreover, the regional pattern of these associations mirrored the regional pattern of associations with BW itself (t_1_ r = 0.57, p_spin_= 0.002; t_2_ r = 0.61, p_spin_= 0.002) (**Supplementary Figure 3**).^38^ Unlike BW itself, PGS-BW did not show a relationship with conditional-longitudinal centiles (**Supplementary Tables 6-8**).

Second, we evaluated a composite score reflecting the burden of neonatal complications to take into account the higher rate of neonatal complications following premature births.^39^ Higher neonatal complications were associated with lower centiles, with strongest effect sizes for subcortical structures at t_1_, with some regions also found to have significant associations at t_2_ (NeoComp_t1_ N_regions_ (p_fdr_<0.05) = 26, β_range_= (−0.056, 0.031); NeoComp_t2_ N_regions_ (p_fdr_<0.05) = 3, β_range_= (−0.048, −0.043)) (**Supplementary Figure 4**). Conditional-longitudinal centiles were not significantly associated with neonatal complications. Findings for BW and GA were largely robust to controlling for neonatal complications (**Supplementary Tables 6-8**).

We conducted sensitivity analyses to confirm the robustness of these findings. First, given the possible relationships between birth-related factors and socioeconomic status, we included income level and parental education level as covariates in the linear regressions and found our results to be largely robust (**Supplementary Figure 5**). Second, given possible relationships between perinatal factors and adolescent body size, we included height and weight as covariates, and similarly found convergent results (**Supplementary Figure 6**). Third, given the possibility that regional effects could be driven by an overarching influence on intracranial size, we included centile scores of intracranial volume as a covariate. Associations with BW and GA for both cross-sectional and longitudinal models were robust to controlling for intracranial volume. However, associations with PGS-BW were no longer significant when controlling for intracranial volume (**Supplementary Figure 7**).

### Increases in psychopathology over time associated with lower conditional-longitudinal brain centiles

Finally, we examined whether conditional-longitudinal centiles, which index change in brain volume between timepoints, were associated with the P-factor, a longitudinally measured behavioral variable. We found that at both baseline and follow-up the P-factor at was negatively associated with cross-sectional centiles, meaning that smaller brain volumes were related to higher psychopathology scores (P-factor_t1_, N_regions_ (p_fdr_<0.05) = 95, β_range_= (−0.080, −0.020); P-factor_t2_, N_regions_ (p_fdr_<0.05) = 81, β_range_= (−0.075, −0.028)) (**Figure 3**).

**Figure 3.**
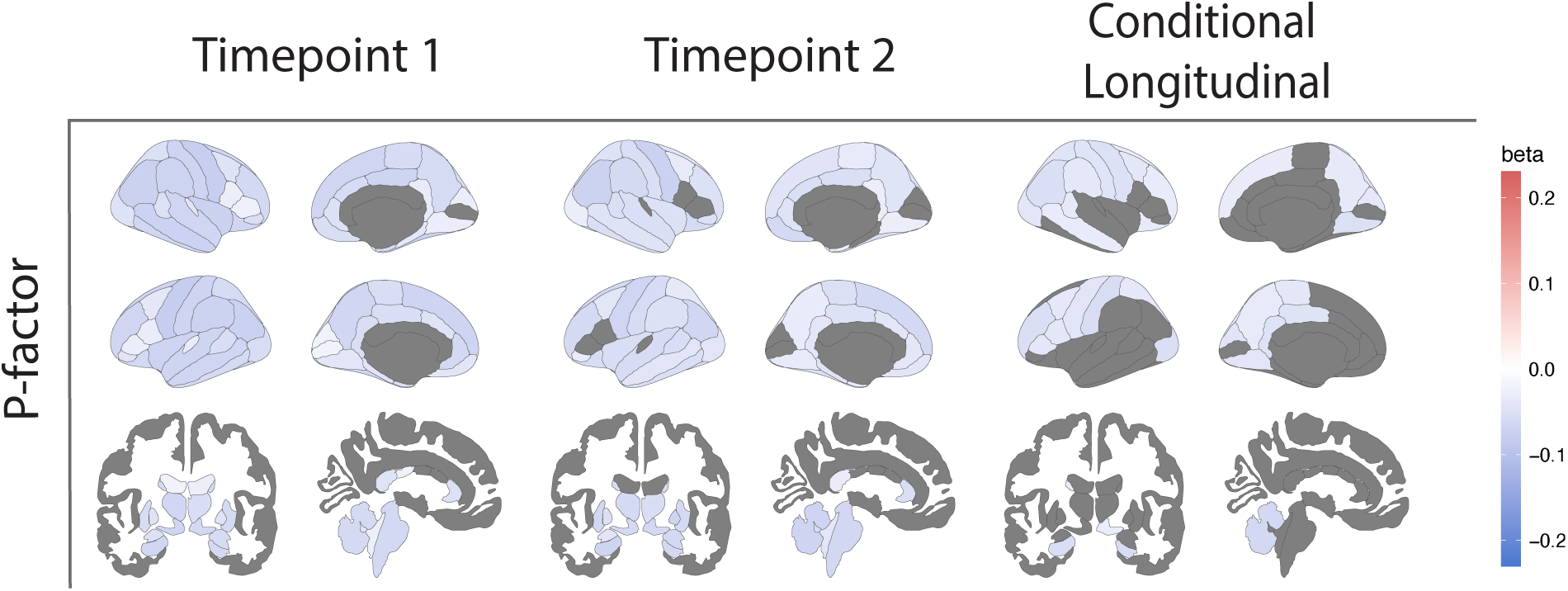
Smaller Longitudinal Centiles Are Associated with Greater Increases in P-factor. Higher psychopathology measured at t_1_ and t_2_ was associated with corresponding lower cross-sectional centiles, and larger increases in psychopathology between timepoints was associated with lower conditional-longitudinal centiles. P-factor_t1_, N_regions_ (p_fdr_<0.05) = 95, β_range_= (−0.080, −0.020); P-factor_t2_, N_regions_ (pfdr<0.05) = 81, β_range_= (−0.075, −0.028); P-factor_long_, N_regions_ (p_fdr_<0.05) = 37, β_range_= (−0.061, −0.031). All results shown are corrected at p_fdr_<0.05, regions that did not survive multiple comparisons are in gray. For detailed information on regional statistics and regions not visualized in this figure see **Supplementary Tables 4-6**.

Next, we tested whether within-individual change in P-factor between timepoints was related to conditional-longitudinal centiles and found a significant negative relationship. Larger increases in P-factor between baseline and follow-up was related to smaller conditional-longitudinal centiles across multiple frontal, parietal, and occipital cortical regions, as well as in the cerebellum and bilateral hippocampus (P-factor _long_, N_regions_ (p_fdr_<0.05) = 37, β_range_= (−0.061, −0.031)) (**Figure 3**). In contrast, within-individual change in P-factor between timepoints showed no significant associations with cross-sectional centiles from either timepoint. This underscores the increased sensitivity of conditional-longitudinal models to person-level changes in longitudinally measured behavioral factors (See **Supplementary Tables 6-8**).

## Discussion

Introducing conditional-longitudinal normative models, we have extended normative modeling used in longitudinal imaging datasets by specifically targeting variability in trajectories of brain development. This framework uniquely allows assessment of individual deviations in developmental trajectories beyond baseline differences in size, which is not feasible with previous cross-sectional normative approaches to longitudinal neuroimaging datasets. In ABCD, we demonstrate that conditional-longitudinal centiles were associated with both perinatal factors and longitudinal changes in psychopathology scores during early adolescence.

Perinatal factors are associated with both cross-sectional variation and longitudinal change in adolescent brain growth. Consistent with previous cross-sectional studies,^17,40–42^ we show that lower cross-sectional centiles, indexing smaller volumes, are associated with lower GA, lower BW, and higher burden of neonatal complications. BW has the strongest associations with cross-sectional brain centiles, and lower BW is also associated with lower conditional-longitudinal centiles in frontal and temporal cortex, bilateral thalamus, and brainstem. Given that most brain regions are decreasing in volume during this age window,^1^ this relationship suggests larger-than-typical declines in regional volumes in participants with lower BWs. Importantly, BW findings are largely robust to controlling for intracranial volume, suggesting localized susceptibility of some brain regions to long-term effects of adverse birth conditions. Associations with BW are largely robust to controlling for GA, socioeconomic status, and adolescent body size—suggesting a relatively specific association between BW and adolescent brain development.

BW is a complex phenotype, related to both underlying genetics and the maternal-fetal environment.^43,44^ Our analysis of PGS-BW provides a new perspective on how these two sources of phenotypic variance could differ in their relationship with future brain development. While we find that PGS-BW is associated with cross-sectional regional volumes at, these associations are not robust to controlling for intracranial volume. Furthermore, PGS-BW is not associated with conditional-longitudinal centiles, suggesting the potential genetic association between lower BW and smaller brain volumes does not extend to the variability in brain maturation dynamics. Importantly, our sample size for PGS-BW is limited to the European-ancestry subsample of ABCD due to ancestry bias in the underlying GWAS which remains unfortunately common in genetic studies.^45^ This results in a smaller sample size which may lead to insufficient power to detect small true effects. Nevertheless, it is plausible that there is a shared genetic basis between BW and intracranial size that is stable over time.^46^ In contrast, the phenotype of BW may have a non-genetic association with dynamic brain changes during early adolescence, due to the contribution of environmental variables that contribute to BW variability. For example, intrauterine conditions leading to lower BW may also lead to epigenetic changes^47^ which continue to influence brain growth through metabolic and other cellular mechanisms.

Our conditional-longitudinal normative models have also shown that within-individual brain changes over time are associated with within-individual changes in psychopathology. These results extend prior reports of cross-sectional associations between brain size and psychopathology in ABCD.^23^ Higher degree of psychopathology at follow-up was associated with smaller than expected brain volumes when accounting for baseline differences in size. Similarly to BW, this relationship was robust to controlling for socioeconomic factors and body size, supporting a relatively specific association between within-individual changes in psychiatric vulnerability and within-individual changes in brain volumes. This association is consistent with prior reports of accelerated thinning and volume reduction correlated with emerging depressive symptoms in smaller samples of adolescents.^25,48^ Greater reduction in brain volumes in adolescents with increasing psychopathology could provide clues into the mechanistic changes in neurodevelopment that confer higher risk for future psychiatric disorders.

We note several methodological issues with the present study. We report on brain volumes within a single cortical parcellation during adolescence. Future work should investigate other morphometric features, alternative parcellations, and studies of the aging brain where conditional-longitudinal centiles may also be particularly useful. While we focus on conditional growth charts rather than velocity growth charts, velocity charts, which are based on the rate of change at fixed time intervals, may be another important application of normative models to longitudinal data.^49^ Finally, while we use publicly released tabular data from ABCD to maximize reproducibility, and follow standard quality control procedures, biases related to suboptimal image quality may still persist in these data.^50^

Notwithstanding these limitations, our findings demonstrate the utility of conditional-longitudinal models to understand inter-individual variability of within-subject change over time. Conditional-longitudinal centiles yield insight into long-debated questions as to how perinatal factors may influence subsequent brain development, and how within-subject brain changes are related to within-subject changes in mental health during adolescence. These models may ultimately inform early identification of adolescents at risk for atypical brain development, providing a framework for preventative interventions.

All data and code are publicly available to facilitate future applications. Our approach may prove useful for investigating deviations from normative trajectories in future clinical and pre-clinical samples, to ascertain how trajectories of psychiatric conditions relate to the dynamics of brain development. Furthermore, the framework of conditional-longitudinal models can be widely applied to longitudinal data from both developing and aging populations, to improve characterization of how the brain continues to change throughout the lifespan.

## Supporting information

STables 3-5

STables 6-8

## Acknowledgments

This work was supported by R01MH132934, R01MH133843, and R01MH134896-01 (PI, AA-B).

## Data Acknowledgement

Data used for analyses presented in this work were obtained from the Adolescent Brain Cognitive Development Study® (ABCD) (https://abcdstudy.org), from the NIMH Data Archive (NDA). Access to the ABCD study data is restricted to researchers with an approved NDA Data Use Certification (DUC).

