## Supplemental Methods, Figures, Table S1 and S2 for "Conditional-longitudinal brain growth charts detect MRI changes with birth weight and psychopathology"

**Affiliations:**

S1. Sample Selection

ABCD Study exclusion criteria cover severe conditions which would interfere with the study protocols[1], as well as MRI scanner contraindications[2]. Participants were excluded due to missing age data, and sex other than male or female. Image quality ratings provided with the ABCD data release were used to filter the sample based on the T1-weighted image inclusion flag. For conditional-longitudinal analyses, participants with different site codes for $t_1$ and $t_2$ were removed to ensure appropriate harmonization of site effects. Detailed numbers on sample selection are outlined below:

From an initial sample of N = 11,868 participants, three participants were removed for no developmental history data, three participants for sex recorded other than male or female, and one participant for missing recorded age at both baseline and timepoint 2.

Next, we filtered based on imaging data and quality control (QC) measures. For $t_1$, 139 participants were excluded for missing imaging data, and 458 participants were excluded due to a "don't include" QC flag, leaving N = 11,264 participants. Seven participants were further excluded for having site IDs that were unmatched in the ABCD Reproducible Matched Samples data split table[3] resulting in 11,257 participants. For $t_2$, 2877 participants were excluded due to missing imaging data, and 196 participants were excluded due to a "don't include" QC flag, leaving 7895 participants. Of these, 333 participants without baseline data were removed, leaving N = 7562 participants.

Data splits using the ABCD Reproducible Matched Samples (ARMS) matched splits[3] were created for split-half model fitting, and additional exclusion were applied to prevent data leakage. At $t_1$, during training, participants with duplicate family IDs between the two split-half sets were removed, with $N_{split\text{-}A\text{-}train}$ = 4535 (N = 886 removed) and $N_{split\text{-}B\text{-}train}$ = 4512 (N = 897 removed). During testing, participants with overlapping family IDs between the two split-half sets were removed, resulting in $N_{split\text{-}A\text{-}test}$ = 5421 (N = 73 removed) and $N_{split\text{-}B\text{-}test}$ = 5409 (N = 68 removed). At $t_2$, we applied the same exclusions as detailed above with a final sample of $N_{split\text{-}A\text{-}test}$ = 3674, $N_{split\text{-}B\text{-}test}$ = 3588), and $N_{split\text{-}A\text{-}train}$ = 3053, $N_{split\text{-}B\text{-}train}$ = 2,962. For longitudinal modeling, participants whose site IDs did not match between $t_1$ and $t_2$ (N= 74) were also removed, with a final sample of $N_{split\text{-}A\text{-}test}$ = 3643, $N_{split\text{-}B\text{-}test}$ = 3549 and $N_{split\text{-}A\text{-}train}$ = 3028, $N_{split\text{-}B\text{-}train}$ = 2929.

#### S2. Imaging-derived phenotypes (IDPs)

*Acquisition and preprocessing*

Volumetric data were derived from T1-weighted images for the ABCD Data Release 5.1[4]. Complete details on processes related to MRI acquisition and pre-processing are described in other reports[2,5]. Whole-brain T1-weighted images were collected with 1mm isotropic voxels and varying parameters based on scanner type. In brief, processing steps included correcting for gradient nonlinearity distortions[6], intensity nonuniformity correction based on tissue segmentation and sparse spatial smoothing, and resampling with 1mm isotropic voxels into rigid alignment with an atlas brain. Freesurfer v5.3.0 was used for cortical surface reconstruction, including skull-stripping[7], white matter segmentation, initial mesh creation[8], correcting topological defects[9,10], surface optimization[8,11,12], and nonlinear registration to spherical surface-based atlas[13]. Cortical regional IDPs were labeled using the Desikan-Killiany atlas-based parcellation[14] and subcortical IDPs were labeled with 'Aseg' atlas-based segmentation[15], which also included cerebellar gray and white matter for left and right hemispheres, as well as six ventricular regions.

Imaging quality control was implemented via the *qc_t1_flag* in the tabulated data, which included inspection by trained technicians for poor image quality, and severe imaging artifacts that may interfere with downstream segmentation[5].

*Global IDPs*

Total cerebral white matter was defined as the sum of left hemisphere and right hemisphere white matter. Subcortical GMV was defined as the sum of the bilateral thalamus, caudate, putamen, pallidum, hippocampus, amygdala, accumbens, and ventral diencephalon. Ventricle volume was defined as the sum of the left lateral ventricle, right lateral ventricle, left inferior lateral ventricle, right inferior lateral ventricle, 3rd ventricle, and 4th ventricle. Intracranial volume was derived from the intracranial volume ROI in the Aseg[15] segmentation in Freesurfer.

S3. Birth-Related Factors

*Questionnaire data*

Gestational age at birth was calculated by subtracting the number of weeks reported premature (derived from the *Q:devhx_12_p* field in the ABCD Data Release) from 40 weeks of gestation. For participants not born prematurely (derived from *Q:devhx_12a_p*), gestational age was encoded as 40 weeks, keeping with previous studies of gestational age in the ABCD dataset[16]. (We also repeated our main analyses with an alternative encoding of term-born subjects as 38 weeks and found that the primary results were largely unaffected (**Supplementary Tables 6-8**).)142 participants had no information on prematurity, and an additional 27 had missing information on "weeks born prematurely", resulting in 169 participants being excluded from gestational age-related analyses. Of note, participants with less than 28 weeks' gestation were excluded from the ABCD study[17].

Birth weight was calculated in ounces, summing pounds reported and ounces reported. For participants with missing ounces data (N = 693), 8 ounces were added to the reported pounds to reflect the most likely average weight and to maximize inclusion in our analyses. 487 participants lacked birth weight in pounds and were excluded from birth-weight–related analyses. Birth weight percentile was calculated using the *growthstandards* package v0.1.6[18] in R, which uses Intergrowth standards[19] birth weight charts to calculate the percentile for birth weight that is based on gestational age at birth and sex.

Neonatal complications were derived as a composite sum of the total number of birth complications endorsed, with a maximum of 8 points. "Don't know" responses (n = 999) were treated as missing. Scores were calculated ignoring missing items unless all responses were missing, resulting in 154 participants with missing overall neonatal complication scores.

Polygenic scores

Saliva samples were collected at the baseline time point of the ABCD study20. DNA was genotyped using custom Smokescreen™ genotype array, containing over 300,000 single nucleotide polymorphisms (SNPs). Prior to imputation, SNPs with a genotyping rate <90^% were removed. Samples with a genotyping rate <95%, samples where genotyped sex and reported sex were mismatched, and samples with excessive heterozygosity were further excluded. To ensure accurate calculation of Hardy-Weinberg Equilibrium (HWE) and heterozygosity, these quality control steps were performed within genetic ancestral groups identified through principal-component (PC) clustering on data combined with the 1000 Genomes Phase 3 data[20]. Genetic ancestry QC was completed as detailed elsewhere[21].

Polygenic score for birth weight ($PGS_{birth\ weight}$) was calculated using PRS-CS[22] using the genome-wide association study (GWAS) from the Early Growth Genetics[23] (EGG ) consortium. PRS-CS is a Bayesian algorithm that infers SNP posterior effect sizes using continuous shrinkage, without the necessity to define p-value thresholds. Following recommendations for highly polygenic traits, global shrinkage prior (phi) was set to $1x10^2$. As the GWAS was conducted in individuals of European ancestry[24], $PGS_{birth\ weight}$ could only be reliably calculated in the in the European-ancestry subsample of ABCD, identified by genetic PCs. For all analyses, all PGS were standardized with a mean of zero and a standard deviation of 1, and the first ten genetic principal components were included as covariates.

S4. P-factor

At baseline and year two of the ABCD Study, youths and caregivers complete mental health questionnaires, including the Kiddie Schedule for Affective Disorders and Schizophrenia (KSADS-5), Prodromal Questionnaire–Brief Child Version (PQ-BC), Child Behavior Checklist (CBCL), and General Behavior Inventory 10 item Mania Scale (GBMI)[25]. To capture overall psychopathology, we used a longitudinal bifactor model in which each questionnaire item loads on a general factor ("P-factor") and one of the orthogonal sub-factors. We used a split-sample exploratory-to-confirmatory procedure similar to Moore et al (2020)[26]. Briefly, data were randomly split into exploratory and confirmatory samples, ensuring that siblings were in the same sample to avoid leakage. In the exploratory sample (N = 5,938, independent measures = 11,119), exploratory structural equation modeling (ESEM) was used for item factor analysis of 178 individual items. The analysis was stratified by site, clustered by family, and grouped by the two timepoints, constraining the model loadings and thresholds to be invariant across time. Model fit was assessed using root mean-square error of approximation (RMSEA; acceptable = 0.06), comparative fit index (CFI; acceptable = 0.95), Tucker-Lewis index (TLI; acceptable = 0.95), and standardized root mean-square residual (SRMR; acceptable = 0.08), where acceptability cut-offs were set to suggestions from Hu and Bentler (1999)[27]. Confirmatory factor analysis (CFA) was performed in the confirmatory subsample. CFA was consistent with the ESEM in site stratification, family clustering, and grouping by time-point. Model fit was assessed as detailed above. Fit was acceptable in both exploratory and confirmatory models, with 125/178 items with loadings >0.40 used in the re-estimated on the full sample. The final bifactor model included loadings onto the general p-factor, as well as 8 orthogonal sub-factors. For analyses presented in this study, we used only the general p-factor scores for both timepoint 1 and timepoint 2. For the general factor, bifactor-associated reliability and generalizability indices were above accepted cutoffs[28]: omegaH = 0.83, H = 0.98, factor determinacy = 0.97, percent uncontaminated correlations = 0.82.

#### S5. GAMLSS Modeling

Global and regional brain volumes from anatomical scans were extracted for $t_1$ and $t_2$. Separate models were fit for 4 global measures of brain volume (Total Cortical Gray Matter, Total White Matter, Subcortical Gray Matter, Total Ventricular Volume), for intracranial volume, and for regional volumes defined by the Desikan-Killiany cortical parcellation, as well as for subcortical structures.

GAMLSS models were fit using a split-half cross-validation approach where splits were based on the ARMS as described in S1[29]. To prevent data leakage due to related individuals, we removed participants with family IDs that repeated between split-halves (randomly removing participants from one of the split halves). For model training, we additionally retained only one random participant from each family.

Parameters of final models are reported in **Supplemental Tables 3-5**.

*Distribution Family Selection*

As the first step of the GAMLSS pipeline, we tested 3 and 4 parameter distributional families in the full sample global IDPs for the cross-sectional $t_1$ and conditional longitudinal $t_2$ models. Fits of distribution families were assessed based on visual inspection of worm plots and comparison of BIC scores. The Box-Cox t-distribution (BCT) was the best-fitting family when considering both cross-sectional and conditional-longitudinal models. For consistency across our models, we thus used BCT as the distribution family for all GAMLSS models. BCT has four parameters ($\mu, \sigma, \nu$,

$\tau), where\ \ \mu > 0,\ \ \sigma > 0,\ \ \nu = (-\infty, \infty)$, and $\tau > 0$. These parameters control the location, shape, and scale of the distribution: $\mu$ is the median, $\sigma\left(\frac{\tau}{\tau-2}\right)^{0.5}$ is approximately the coefficient of variation, $\nu$ controls the skewness, and $\tau$ controls the kurtosis[30,31].

*Model Fitting*

Model fitting procedures were identical for cross-sectional models in $t_1$ and $t_2$. Model selection space included a linear effect of age and a fixed effect of sex as predictor variables for *mu*, *sigma*, and *nu* and a constant for *tau*. We additionally tested an interaction for age and sex in *mu*. We assessed nonlinear age effects, but linear age effects were found to be optimal for all models, likely due to the relatively narrow age range. All models also included a random effect of site in *mu* and sigma to account for site-specific effects in the dataset, which in practice has been shown to result in similar harmonization as ComBat[32].

Model selection was conducted in parallel for each split-half. For each IDP, the model with the lowest BIC was selected (**Supplemental Tables 3-4**). Fits of the final models were further assessed via correlations between expected and actual proportion of subjects below a given centile (min $r > 0.999$).

*Conditional Longitudinal Models*

Brain volume centiles derived from cross-sectional models are informative in assessing where an individual may fall with regards to the distribution of the population. However, a strictly cross-sectional approach to brain volumes misses how these volumes change over time, which can also differ substantially between individuals. To model longitudinal measurements of brain volumes in our data, we fit GAMLSS models to $t_2$ volumes, conditioned on $t_1$ volumes for the location parameter (*mu*). Model selection was conducted similarly to cross-sectional models, with the interscan interval between $t_1$ and $t_2$ added as a possible covariate for *mu*. Interscan interval was calculated as months between recorded age at $t_2$ and recorded age at $t_1$(**Supplemental Table 5**).

*Centiles*

Individual centiles are relative to the reference normative model, chosen through the model selection processes detailed above. For conditional longitudinal models, individual centiles are based on the same individual's $t_1$ measurements, providing a measure of deviation for that individual's $t_2$ measurement, given their $t_1$ measurement. For each IDP, we calculated cross-sectional and conditional-longitudinal centiles for each subject in the dataset. Centiles were calculated for each split-half test set, using the model fitted to the other split-half training set. As an additional quality check, we calculated correlations between centiles using the model fitted to the same vs. other split-half and found a high degree of correspondence across all models (min $r = 0.971$).

*Centile Mahalanobis Distance*

As a cumulative measure of normative deviation across all centiles[32], we calculated the Centile Mahalanobis distance (CMD)[33] using centiles for 100 cortical and subcortical regional volumes. CMD is formalized as:

$$D_M(x) = \sqrt{(\mathrm{x} - \mu)^T S^{-1} (\mathrm{x} - \mu)}$$

where $x$ denotes a set of observations across multiple phenotypes, $\mu$ denotes the mean across those observations, and $S$ denotes the covariance matrix across both. Squared Mahalanobis distance is equal to the sum of squares of non-zero standardized principal component scores. Thus, CMD is a metric of the distance between an individual's centiles and the center of the normative multi-phenotype (multi-dimensional) centile space, taking into account the correlated structure of the multiple phenotypes. The CMD was used to compare an individual participant's set of centiles across the cross-sectional and conditional longitudinal models. Relationships between CMDs and regional centiles between $t_1$, $t_2$, and conditional longitudinal models were assessed using Pearson's correlation.

#### S6. Additional Covariates in Sensitivity Analyses

*Income and Parental Education*

We derived income and parental education from parental-report questionnaires[25]. Parental education categories were condensed to "Didn't complete high school", "Completed highschool/GED”, “Some college/Associate's Degree”, “Bachelor's Degree”, and “Advanced Degree” to assure appropriate distribution across the factor levels. For subjects missing income or parental education responses at $t_1$, we used responses from $t_2$ if available. In linear regressions, income and parental education were encoded as ordinal variables.

*Weight and Height*

Weight and height measures were derived from anthropometric data collected by the ABCD Study at both timepoints[25]. Recorded averages of three repeated measurements taken at each timepoint were included as covariates for the analyses involving $t_1$ and $t_2$ models respectively. For conditional longitudinal models, the measurements from $t_2$ were included as covariates.

*Non-singleton birth status*

Given that non-singleton births are more likely to be born premature[34], we also assessed singleton birth status as an additional covariate in the model selection space of normative models, finding that all main results were robust to this sensitivity analysis.

#### Supplementary Figures

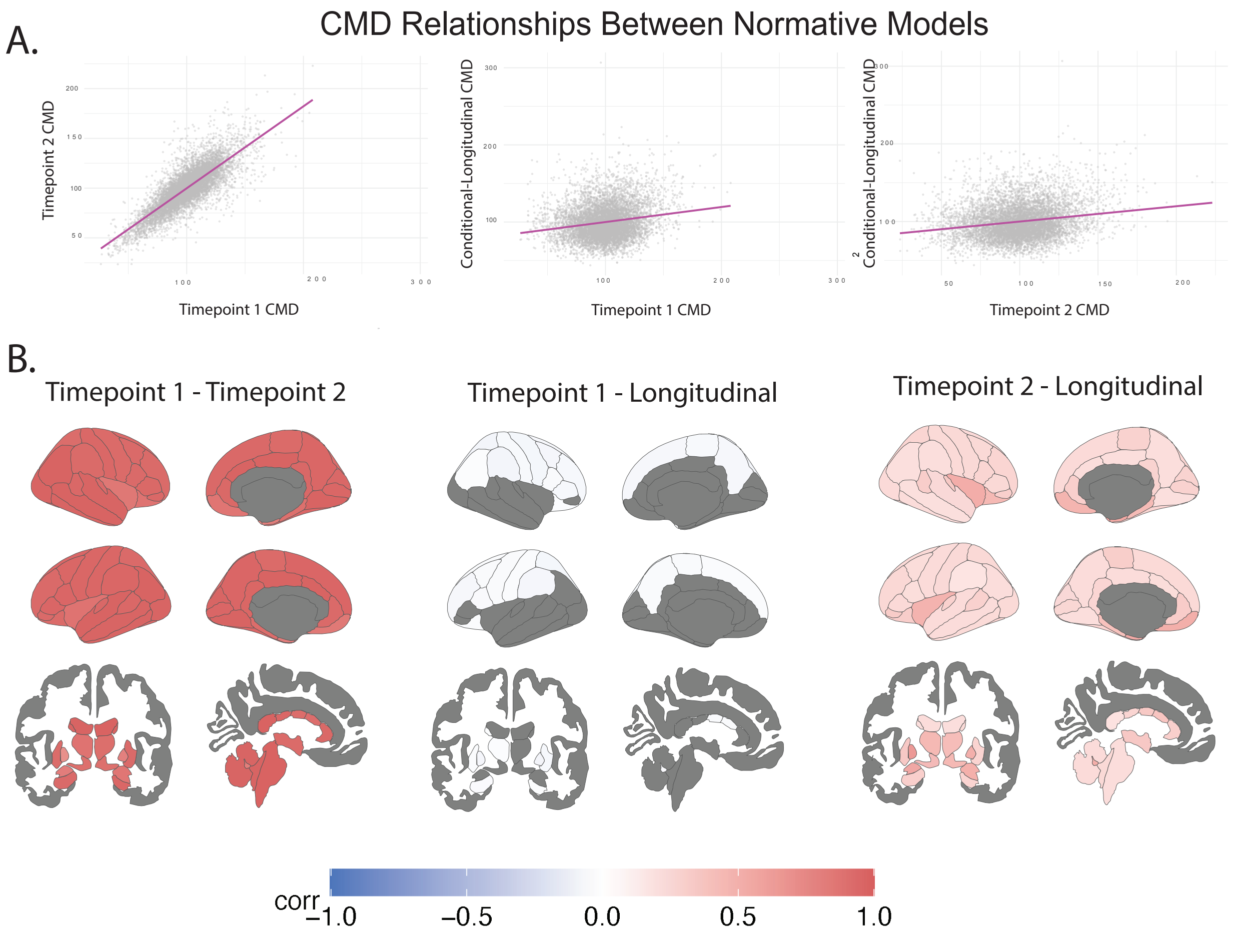


*Supplementary Figure 1. Conditional longitudinal centiles are related to, but distinct from cross-sectional centiles. (A) CMD, a cumulative measure of individual deviations across brain regions is highly correlated between cross-sectional centiles from $t_1$ and $t_2$ ($r = 0.81$, $p < 2.2e\text{-}16$), while the correlation between conditional longitudinal CMD and cross-sectional CMDs are smaller ($corr_{t1} = 0.18$, $corr_{t2} = 0.19$, $p < 2.2e\text{-}16$). (B) Individual deviations for each brain region show similarly high degree of correlation between centiles from $t_1$ and $t_2$ (max $r = 0.99$, min $r = 0.71$). Correlations between conditional longitudinal and cross-sectional centiles show patterns of weak negative correlations for $t_1$(max $r = -0.095$, min $r = -0.030$), and weak to moderate positive correlations for $t_2$(max $r = 0.67$, min $r = 0.13$).*

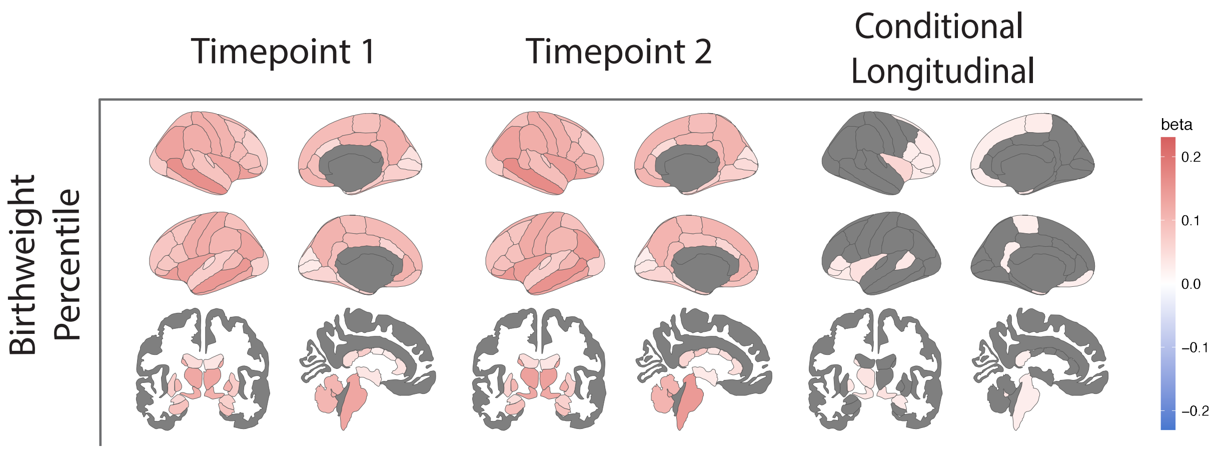


*Supplementary Figure 2. Birth weight percentile (birth weight-for-gestational-age) shows lower strength but similar pattern of associations with centiles in both cross-sectional and conditional longitudinal models compared to birth weight before correction. BWpercentile$_{t1}$: no. regions = 98, $\beta_{range}$= (0.018, 0.16); BW-percentile $_{t2}$: no. regions = 96, $\beta_{range}$= (0.022, 0.17); BW-percentile $_{long}$: no. regions = 31, $\beta_{range}$= (0.023, 0.59). All results shown are corrected at $p_{fdr}$<0.05. Beta scale from -0.23 to +0.23. For detailed information on regression statistics across all regions and global tissue volumes see **Supplementary Tables 6-8**.*

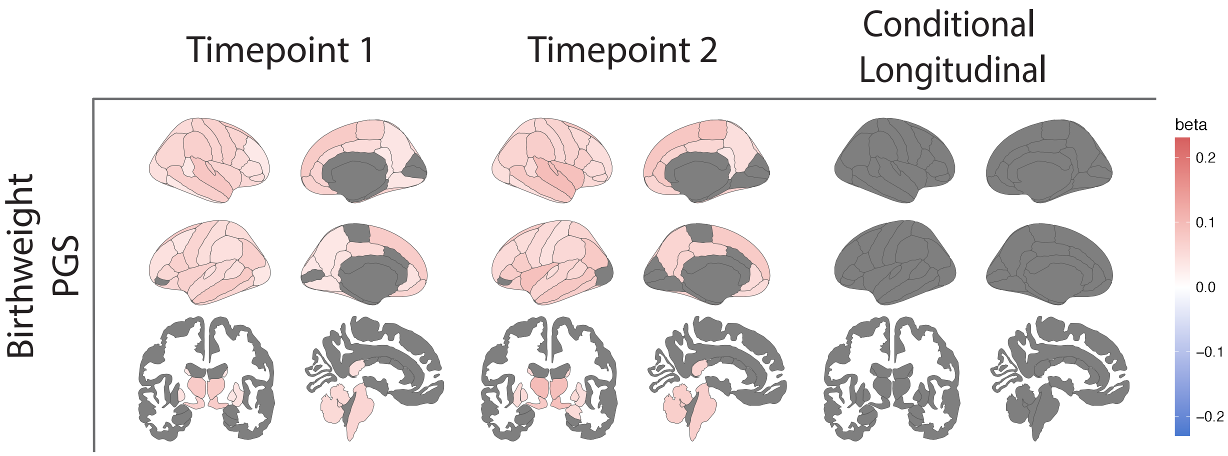


*Supplementary Figure 3. Polygenic score for birth weight ($PGS_{bw}$) shows similar pattern of regional associations with cross-sectional centiles as the phenotype of birth weight but is not associated with longitudinal change. PGS-$BW_{t1}$: no. regions = 73, $\beta_{range}$= (0.031, 0.082); PGS-$BW_{t2}$: no. regions = 71, $\beta_{range}$= (0.035, 0.097). No significant associations for conditional longitudinal centiles. All results shown are corrected at $p_{fdr}<0.05$. Beta scale from -0.23 to +0.23. For detailed information on regression statistics across all regions and global tissue volumes see **Supplementary Tables 6-8**.*

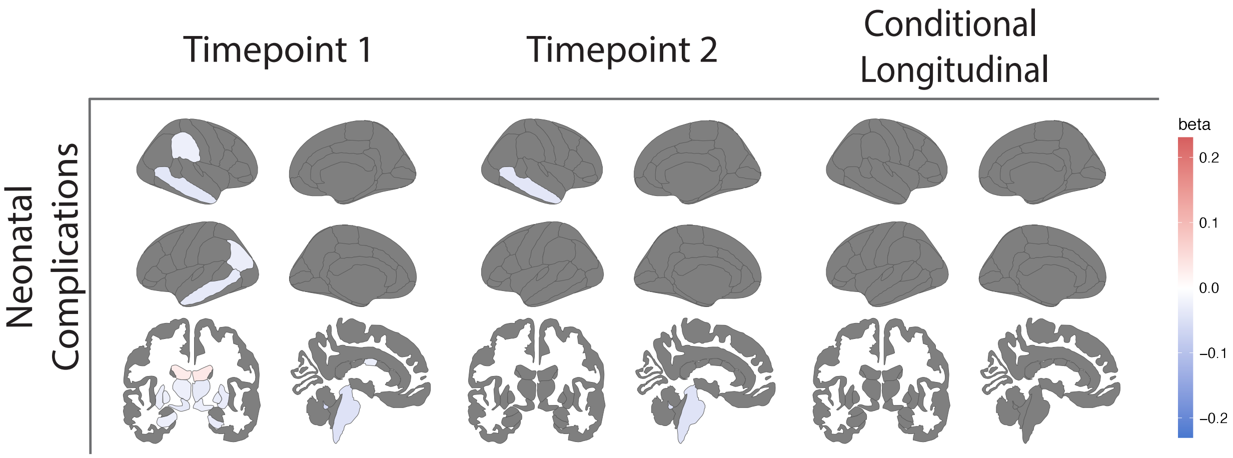


*Supplementary Figure 4. Higher burden of neonatal complications is associated with smaller centiles at $t_1$ mostly in subcortical structures, some temporal and parietal regions, and associated with higher centiles for the lateral ventricles. $NeoComp_{t1}$ : no. regions = 22, $\beta_{range}$= (-0.059, 0.033); $NeoComp_{t2}$ : no. regions = 4, $\beta_{range}$= (-0.049, -0.040). No significant associations for conditional longitudinal centiles. All results shown are corrected at $p_{fdr}$<0.05. Beta scale from -0.23 to +0.23. For detailed information on regression statistics across all regions and global tissue volumes see* ***Supplementary Tables 6-8***.

### Results with Socioeconomic Covariates

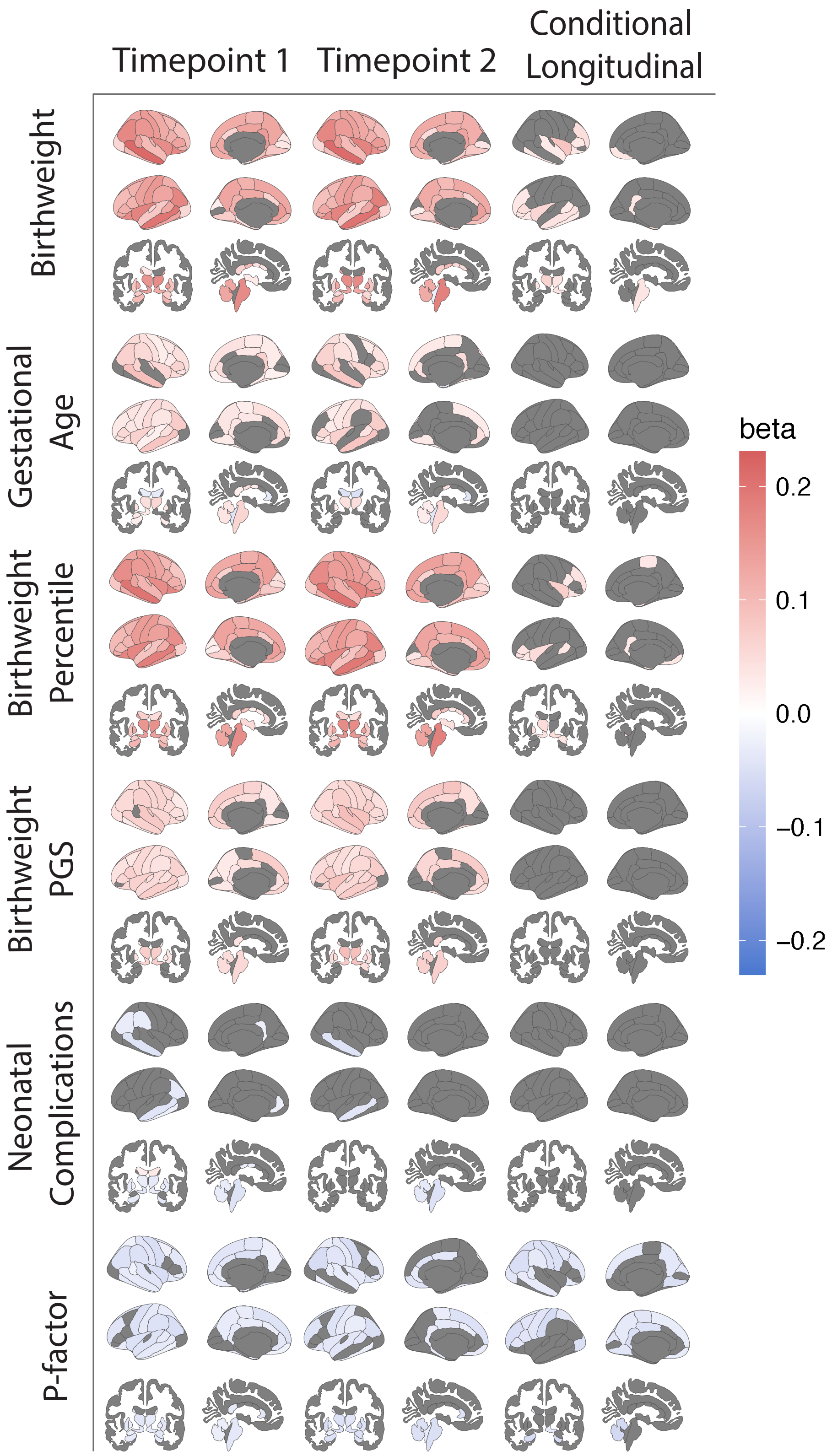


*Supplementary Figure 5. Results are largely robust to controlling for socioeconomic factors. Shown are the standardized beta coefficients from linear regressions where parental income and parental education are included as covariates. All results shown are corrected at $p_{fdr}<0.05$. Beta scale from -0.23 to +0.23. For detailed information on regression statistics across all regions and global tissue volumes see* ***Supplementary Tables 6-8***.

### Results with Weight/Height Covariates

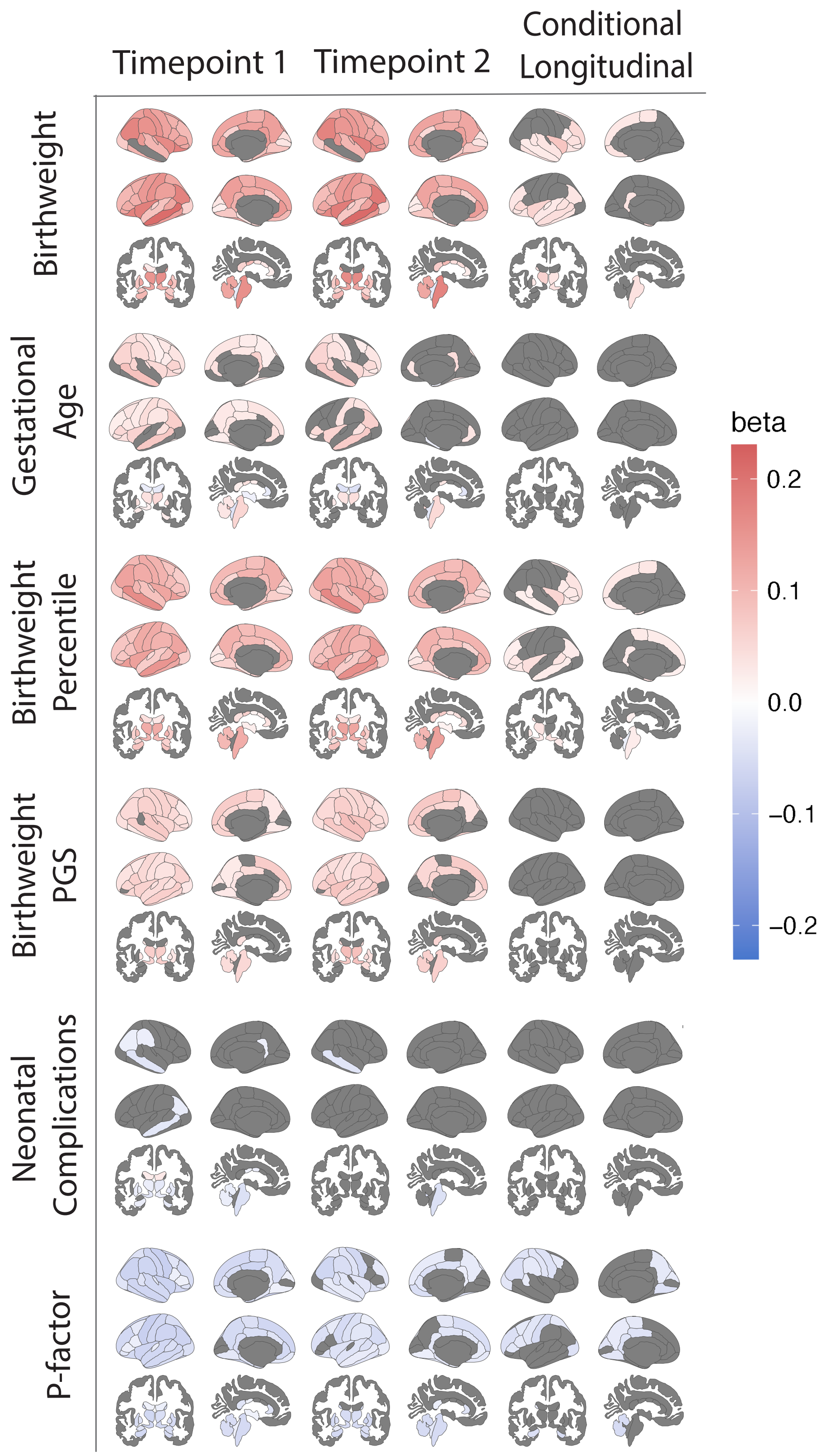


*Supplementary Figure 6. Results are largely robust to controlling for anthropometric factors. Shown are the standardized beta coefficients from linear regressions with adolescent weight and height are included as covariates. All results shown are corrected at $p_{fdr}$<0.05. Beta scale from -0.23 to +0.23. For detailed information on regression statistics across all regions and global tissue volumes see **Supplementary Tables 6-8**.*

### Results Controlling for Intracranial Volume

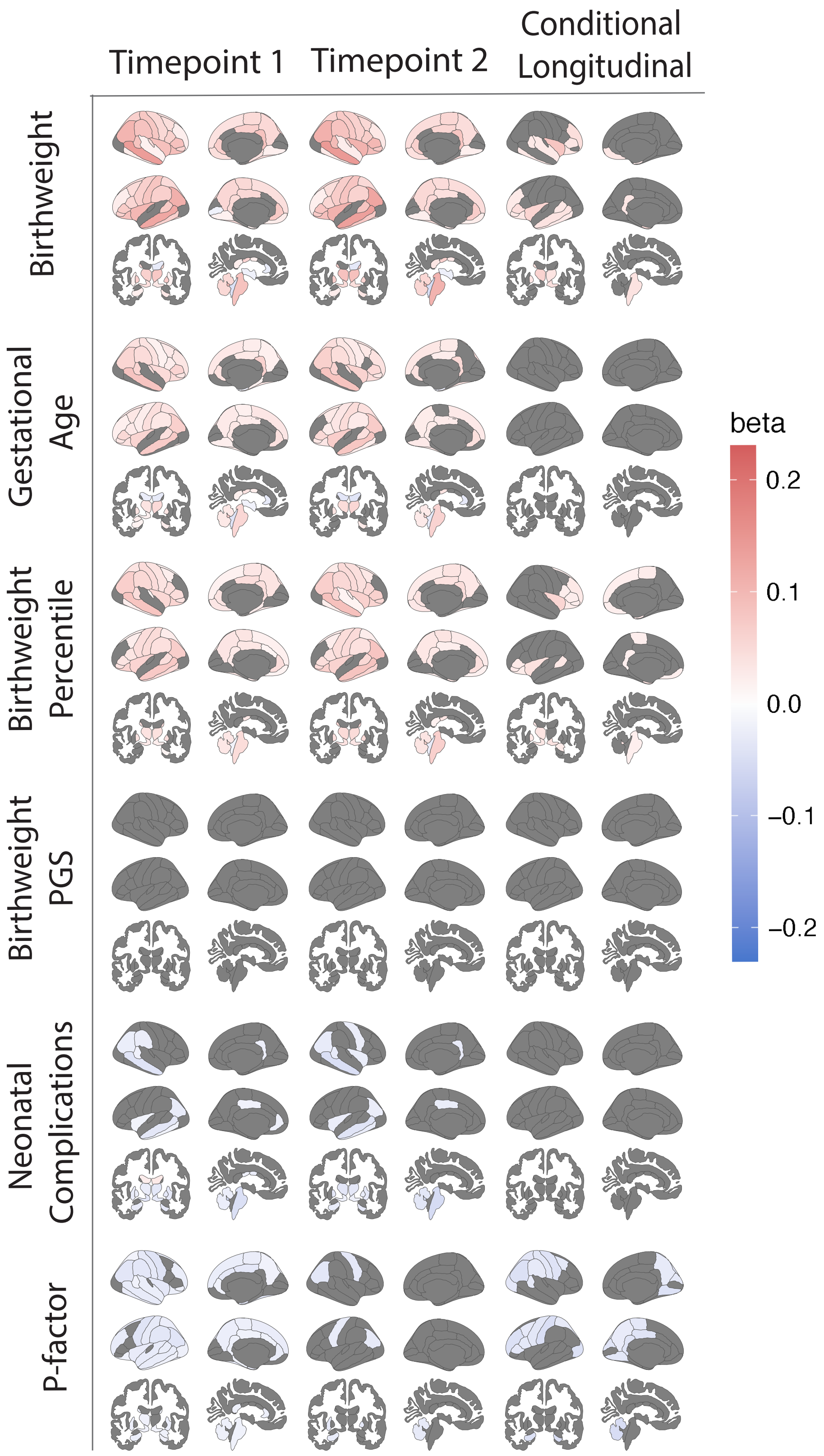


*Supplementary Figure 7. Results for birth weight, gestational age, neonatal complications, and p-factor are largely robust to controlling for intracranial volume. Shown are the standardized beta coefficients from linear regressions with normalized individual centile score for total brain volume is included as a covariate. All results shown are corrected at $p_{fdr}<0.05$. Beta scale from -0.23 to +0.23. For detailed information on regression statistics across all regions and global tissue volumes see **Supplementary Tables 6-8**.*

**Supplementary Tables**

| Question Code | Did he/she have any of the following complications at birth? | Yes n | Yes Nn(before splits) | 999 (don't know n) | 999 (don't know n) (before split-sample) | No Answer n |
|---|---|---|---|---|---|---|
| devhx_14a3_p | Blue at birth? | 341 | 352 | 293 | 300 | 1 |
| devhx_14b3_p | Slow heartbeat? | 300 | 305 | 310 | 318 | 1 |
| devhx_14c3_p | Did not breathe at first? | 508 | 520 | 267 | 277 | 1 |
| devhx_14d3_p | Convulsions? | 16 | 17 | 199 | 207 | 1 |
| devhx_14e3_p | Jaundice needing treatment? | 1720 | 1782 | 260 | 269 | 1 |
| devhx_14f3_p | Required oxygen? | 1047 | 1071 | 267 | 273 | 1 |
| devhx_14g3_p | Required blood transfusion? | 54 | 56 | 185 | 195 | 1 |
| devhx_14h3_p | Rh incompatibility? | 266 | 278 | | 382 | 2 |

Supplementary Table 1. ABCD Developmental Questions for deriving the neonatal complication burden measure.

| Variable | Split half A, t1 Total (n = 5,421) | Split Half B, t1 Total (n = 5,409) | Split half A, t2 Total (n = 3,674) | Split half B, t2 Total (n = 3,588) |
|---|---|---|---|---|
| Weight, mean (SD), lbs | 82 (23) | 83 ( 24) | 108 (32) | 108 (32) |
| Weight, available n | 5416 | 5399 | 3664 | 3579 |
| Height, mean (SD), inches | 55.3 (3.2) | 55.2 (3.3) | 60.2 (3.5) | 60.2 (3.5) |
| Height available n | 5417 | 5403 | 3667 | 3582 |
| Total Household Income, n (%) | | | | |
| Less than $5,000 | 206 (4.0%) | 216 (4.2%) | 112 (3.2%) | 127 (3.6%) |
| $5,000 through $11,999 | 217 (4.2%) | 195 (3.8%) | 131 (3.7%) | 118 (3.4%) |
| $12,000 through $15,999 | 130 (2.5%) | 139 (2.7%) | 71 (2.0%) | 84 (2.4%) |
| $16,000 through $24,999 | 250 (4.8%) | 256 (4.9%) | 170 (4.8%) | 162 (4.6%) |
| $25,000 through $34,999 | 319 (6.1%) | 336 (6.5%) | 216 (6.1%) | 216 (6.2%) |
| $35,000 through $49,999 | 438 (8.4%) | 448 (8.6%) | 327 (9.2%) | 306 (8.8%) |
| $50,000 through $74,999 | 708 (14%) | 726 (14%) | 502 (14%) | 530 (15%) |
| $75,000 through $99,999 | 760 (15%) | 732 (14%) | 541 (15%) | 522 (15%) |
| $100,000 through $199,999 | 1,565 (30%) | 1,557 (30%) | 1,096 (31%) | 1,037 (30%) |
| $200,000 and greater | 600 (12%) | 590 (11%) | 386 (11%) | 392 (11%) |
| Household Income, available n | 5,193 | 5,195 | 3,552 | 3,494 |
| Parental Education, n (%) | | | | |
| Didn't complete highschool | 357 (6.6%) | 355 (6.6%) | 229 (6.2%) | 209 (5.8%) |
| Completed highschool/GED | 561 (10%) | 570 (11%) | 341 (9.3%) | 344 (9.6%) |
| Some college/Associate's | 1,576 (29%) | 1,621 (30%) | 1,071 (29%) | 1,099 (31%) |
| Bachelor's Degree | 1,561 (29%) | 1,480 (27%) | 1,093 (30%) | 1,048 (29%) |
| Advanced Degree | 1,364 (25%) | 1,383 (26%) | 939 (26%) | 888 (25%) |
| Parental Education, available n | 5,419 | 5,409 | 3,673 | 3,588 |

Supplementary Table 2. Distribution of covariates for sensitivity analyses in each split sample and timepoint. SD, Standard Deviation; N, sample size; lbs, pounds. *N available for models including each variable after excluding participants with missing data and exclusions as described in the S6.

Supplemental Tables 3-8 are included in Supplemental Files STable3-5_modelparameters.xlsx, and Stable6-8_regressionstats.xlsx due to their size.
